# Invasive in the North: New latitudinal record for Argentine ants in Europe

**DOI:** 10.1101/837583

**Authors:** N. Pierre Charrier, Caroline Hervet, Claire Bonsergent, Matthieu Charrier, Laurence Malandrin, Bernard Kaufmann, Jérôme M. W. Gippet

**Author notes:** Corresponding authors: NP Charrier, JMW Gippet.

## Abstract

Environmental niche models predict the presence of the invasive Argentine ant in north-western Europe, especially along all the French Atlantic coast. Yet, the species has never been observed North from the 45^th^ parallel in Europe, suggesting either that current models are wrong or that Argentine ants are already spreading north inconspicuously. Here, we report a three-hectare wide colony of Argentine ants, detected in 2016 in Nantes, France, which is 300 km north of the former northern-most outdoor population of this species in Europe. COI sequencing revealed that the haplotype of this new colony is the same as the one found in the so-called Catalonian supercolony, which is distinct from the haplotype found over most of the species range in Europe. Our discovery confirms models’ predictions that Argentine ants can colonize north-western Europe and suggests that they might have already reached several other locations along the French Atlantic coast. Detection surveys should be conducted in order to assess Argentine ants’ invasion patterns in Western France, particularly in high introduction risk areas such as major cities and maritime ports.

## Introduction

The Argentine ant, *Linepithema humile* (Mayr), is a widespread invasive species with strong negative impacts on native biodiversity and human activities (Cole et al. 1992, Suarez et al. 2005, Menke et al. 2018). The species originates from subtropical South America and was unintentionally introduced by human activities on all continents and several islands (Wetterer et al. 2009, Janicki et al. 2016). In Europe, one vast and one smaller supercolonies (i.e. groups of colonies or populations which exhibit no inter-aggressive behavior) which originated from two distinct introduction events, are spread all over the Mediterranean coast and the Iberic peninsula (Giraud et al. 2002). The smallest supercolony was found near Barcelona and has thus been named the Catalonian supercolony (Giraud et al., 2002), while the largest has been named the Main European supercolony. Environmental niche models predict that most Western Europe is already suitable for the establishment of *L. humile*, even at northern locations such as Brittany (France) or England (Roura-Pascual et al. 2004, Bertelsmeier and Courchamps 2014). Yet, Argentine ant populations have never been detected in northern Europe so far (except for indoor introduced colonies; Gómez et al. 2005, Janicki et al. 2016, Blatrix et al. 2018). Considering the ease with which this species is dispersed by humans, it is unlikely that dispersal limitations are responsible for this pattern. Therefore, this unexpected absence could result from two different types of gaps in our understanding of spread in Argentine ants. First, predictive models may be misleading, and the North of Europe is climatically unsuitable for Argentine ants. Second, the species may have been overlooked and is currently spreading northward inconspicuously. Any information that could help identify which of these two hypotheses is the most plausible is therefore of great value for designing either better predictive models or detection surveys that would help eradicate new infestations and prevent further unintentional human-mediated dispersal events (Ujiyama and Tsuji 2018).

Here, we report and document the northernmost colony of Argentine ants occurring outside buildings in Europe. The colony was detected accidentally in 2016 in the urban area of Nantes, France, approximately 300 kilometers north from its outdoor northernmost previously known location in Bordeaux (inpn-mnhn.fr, Blatrix et al. 2018, Janicki et al. 2016). The surface occupied by the colony was measured and mitochondrial DNA (Cytochrome Oxidase subunit I) was sequenced in order to identify the genetic origin of this new population.

## Material and methods

### Detection and colony measurement

Following the first detection in May 2016, the colony was visited in May 2017 and measured on May 8^th^, 2018. The surface area occupied by the colony was measured by a team of three persons searching for nests and trails on the ground, trees and shrubs in every direction from the initial point of detection. The presence of *L. humile* nests or trails was assessed, and workers were collected, approximately every 40 meters or until colony boundaries were reached. All observations (of *L. humile* and native species) and samples were precisely georeferenced. Colony boundaries were reached when, in every direction, no *L. humile* workers were found for more than 100 meters from the closest nest or trail. All mapped *L. humile* occurrences were then imported into ArcGIS v.10.1 (ESRI, 2012) and the surface area occupied by the colony was calculated as the area of the polygon obtained from mapped occurrences by minimum bounding geometry.

### Molecular identification

Argentine ant workers were collected at the detection site in 2017 and preserved in 96°C alcohol. One ant was cut in two parts and its genomic DNA was extracted using a non-destructive protocol with ammonium hydroxide 0.7M. A fragment of 850 bp of the Cytochrome Oxidase subunit I (COI) of the ant was amplified by polymerase chain reaction. Five μL of DNA were added to 25 μL of a mix containing 1 X GoTaq buffer (Promega), 4 mM MgCl_2_, 0,2 mM each dNTP (Eurobio), 1 unit GoTaq G2 Flexi DNA Polymerase (Promega) and 0,5 μM of each primer (see below). Cycling parameters were: 95°C for 5 min followed by 40 cycles of 95°C for 30 sec, 60°C for 30 sec, 72°C for 1 min and 15 sec, and a final extension at 72°C for 5 min. Primers LH01_F (AGGAGCCCCAGATATAGCAT) and LH01_R (GGTATCATGAAGAACAATGTCAA) were designed using Primer3 software (Untergasser et al. 2012) based on the COI coding sequence (GenBank accession number: KX146468.1). Amplified fragments were purified with ExoSAP-IT reagent following manufacturer’s recommendation (Affymetrix) and bidirectional Sanger sequencing was conducted by Eurofins company using the same primers. The COI sequence obtained was aligned alongside 22 other COI sequences belonging to native and introduced populations of Argentine ants from different geographic origins (Vogel et al. 2010, Inoue et al. 2013). Seven COI sequences from closely related *Linepithema* species (*L. oblongum* and *L. gallardoi;* Wild 2009) were used as outgroup. Sequences were aligned using clustalΩ and visualized in Seaview (Gouy et al. 2009). Phylogeny was inferred using a maximum likelihood method with IQ-TREE (Nguyen et al. 2015) under the best model (TIM2+F+G4) found by Model Finder (Kalyaanamoorthy et al. 2017) with Subtree Pruning Regrafting and Nearest Neighbor Interchange (SPR & NNI).

## Results

### Detection and colony measurement

*Linepithema humile* was first detected, accidentally, in May 2016 in the *Ile de Nantes*, an island located on the Loire River in the center of the city of Nantes, France (see Fig. 1a, b). The colony was detected in the western part of the island, on the site of the *Machines de l’île*, a formerly industrial area (shipyard from 1760 to 1987) and currently a commercial and touristic area (second biggest wholesaling platform in France; Fig. 1b, c). The colony was successfully found again in May 2017 and May 2018. During colony measurement, nests were found in the ground or in cracks in the pavement and trails mainly along sidewalks. The surface area of the colony of *L. humile* was 3.1 hectares (30,975 square meters; Fig. 1c). Only one native ant species, *Plagiolepis pygmaea*, was observed inside the invaded area. Other native species (*Lasius niger, Lasius emarginatus, Tetramorium* sp. and *Formica rufibarbis*) were found only outside the invaded area.

**Fig. 1.**
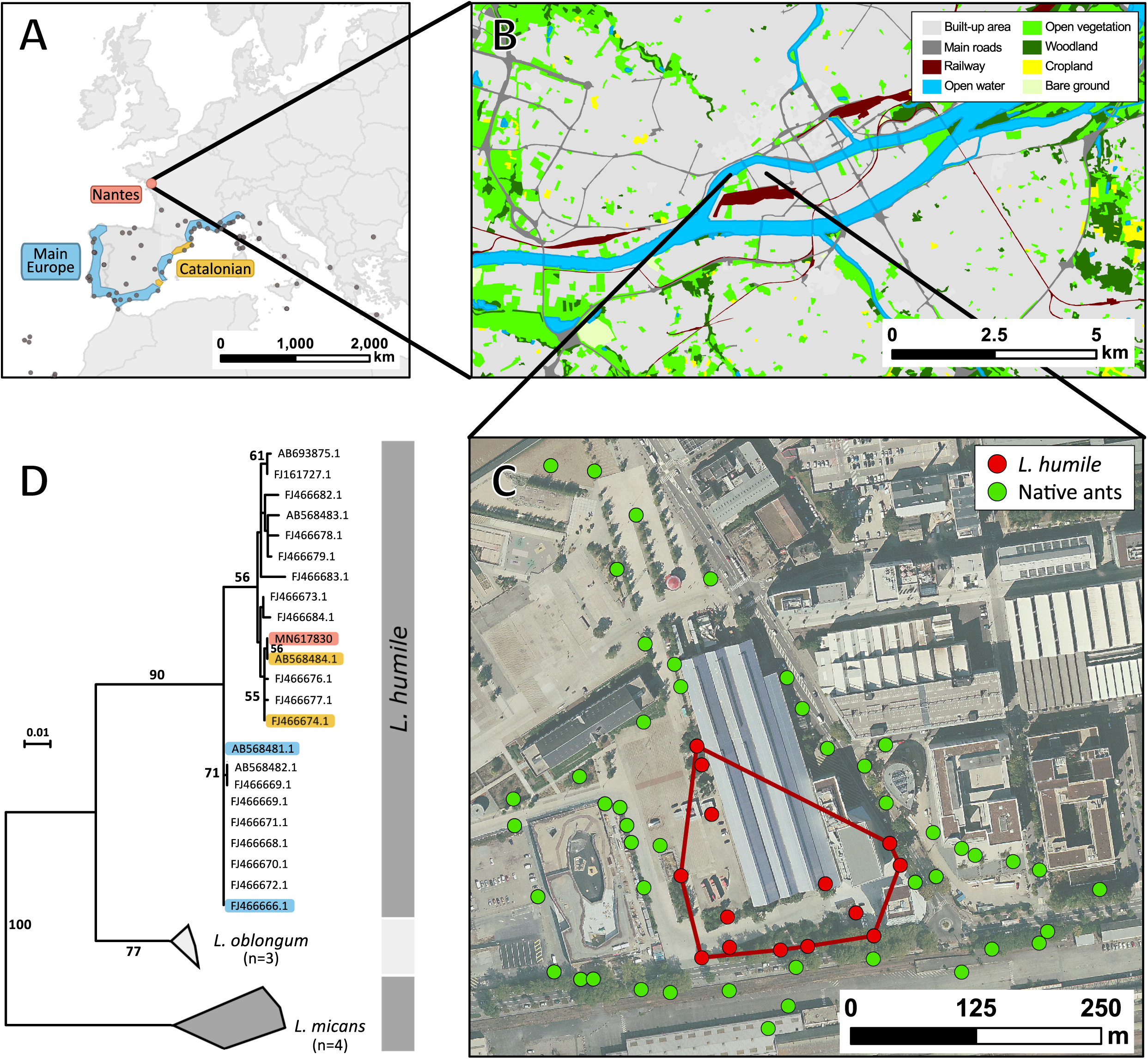
Geographic and genetic situation of the Nantes colony of *Linepithema humile*. **a** Localization of the Nantes colony and other recorded outdoor invaded localities (grey dots) according to AntWeb (antweb.org), AntMaps (antmaps.org) and INPN-MNHN (inpn.mnhn.fr) databases. Colored areas correspond to the position of the two European supercolonies according to Giraud et al. 2002. **b** Landscape situation of the Nantes colony (Background map: data.nantes.fr). **c** Local situation of the Nantes colony (Background map: vuduciel.loire-atlantique.fr) and position of the invasive and native ants’ nests and trails. **d** Phylogenetical placement of the Nantes colony (LH01_cons2) according to COI sequence alignment with 29 other sequences from *L. humile* and two other *Linepithema* species. Bootstrap values are represented on branches leading to a node supported by more than 50 non-parametric bootstrap replicates. Blue and orange labels represent sequences sampled respectively from the Main European and the Catalonian supercolonies (see Fig. 1a).

### Molecular identification

Using *Linepithema humile* (LH) primers, we obtained a 815 bp long DNA sequence corresponding to the COI fragment – sequence deposited under the genbank accession number MN 617830. This sequence was aligned with 29 other COI fragments obtained from previous studies and resulted in an alignment of 741 nucleotides among which 102 were informative. The alignment was used to infer a phylogenetic tree, represented in Fig. 1d. Our sequence was identical to the sequence AB568484.1 (100% identity all along their 749 bp common sequence) which belongs to the Catalonian supercolony (orange highlighted in Fig. 1d; this sequence corresponds to the LH6 haplotype in Inoue et al. 2013). The second sequence available for the Catalonian supercolony (FJ466674.1, haplotype H9 in Vogel et al. 2010) differed from Nantes’ sequence by one site among their 803 bp long common sequence (>99.9% identity). Sequences for the main European supercolony (blue highlighted in Fig. 1d) differed from Nantes’ sequence by 12 and 13 variable sites over 749 and 803 sites (identity of 98.4%) respectively for AB568481.1 (LH1 in Inoue et al. 2013) and FJ466666.1 (H1 haplotype in Vogel et al. 2010).

## Discussion

We reported the northernmost colony of the invasive ant *Linepithema humile* ever detected outside buildings in Europe (Janicki et al. 2016). The colony was observed three years in a row (in 2016, 2017 and 2018) and covers more than three hectares in the center of Nantes, France. It genetically belongs to the Catalonian supercolony and therefore constitutes the first record of this genetic group in France.

The colony detected in Nantes seems well-established since it was easily found three consecutive years and covers more than three hectares. Its outdoor status is clear as the only building surrounded by the colony is a former shipyard facility, open to the outside (no walls), that cannot provide a heated refuge during winter. The presence of *L. humile* along the French Atlantic coast was predicted by environmental niche models based on climate and land cover information (Roura-pascual et al. 2004, Bertelsmeier and Courchamps 2014). In addition, Nantes is the sixth largest city and the fifth more active maritime port of France (INSEE 2013, statistiques.developpement-durable.gouv.fr) and was identified as a highly suitable port of entry for Argentine ants in western Europe (Bertelsmeier and Courchamps 2014). France has 14 maritime ports along its Atlantic coast and the English Channel (statistiques.developpement-durable.gouv.fr), it is therefore likely that Nantes, rather than an isolated case, is only the tip of a larger invasion. The recent detection of *L. humile* in Bordeaux, another large city on the French Atlantic coast (i.e. 300 km South from Nantes, the former northernmost outdoor location of Argentine ants in Europe; Blatrix et al. 2018), tends to support this hypothesis. *Plagiolepis pygmaea* was the only native ant species found inside the area covered by Argentine ants. Interestingly, the species formed large foraging trails on the ground and trees, climbing up to 2.5-3 meters high to reach aphids (J. Gippet, *pers. obs.*). *P. pygmaea* was already found to co-occur with *L. humile* in Spain (Oliveras et al. 2005), presumably because of its submissive behavior (Abril et al. 2009). Our observation suggests that this pattern might be consistent across the distribution of *L. humile* in Europe.

Surprisingly, Cytochrome Oxydase I (COI) gene sequence revealed that the Nantes colony does not belong to the main European supercolony occurring all along the Mediterranean coast (Fig. 1d), but is genetically identical to the Catalonian colony from Sant Cugat del Vallès, in the suburbs of Barcelona, Spain (LH6 in Inoue et al. 2013). Considering that the Main European haplotype (respectively H1 and LH1 in Vogel et al. 2010 and Inoue et al. 2013) is, by far, the most widespread at both European and global scale (Vogel et al. 2010, Inoue et al. 2013), we would have expected it to be the origin of new populations along the French Atlantic coast (i.e. bridgehead effect; Bertelsmeier et al. 2018). Whether or not the Nantes colony is directly connected to the Catalonian supercolony by human-mediated dispersal could be assessed by further investigating the genetic profile of these populations using, for example, microsatellite or genomic data (Javal et al. 2018). More than a new latitudinal record for the species in Europe, the Nantes colony also constitutes the first record of a second Argentine ant haplotype in France. This result suggests a more complex European invasion history than previously thought (Giraud et al. 2002, Vogel et al. 2010) and should stimulate invasion biologists to genetically identify recently discovered outdoors and indoors populations of Argentine ants in Europe in order to better disentangle its spread over the last decades.

Our findings support the predictions of climatic suitability of north-western France for *L. humile* (Roura-pascual et al. 2004, Bertelsmeier and Courchamps 2014). Early detection is the best strategy to limit the spread of invasive ants because eradicating invasive populations becomes less likely and more expensive as colonies grow large (Ujiyama and Tsuji 2018). Detection surveys might therefore be imperative in order to assess the presence of Argentine ant colonies in strategic locations along the French Atlantic coast, such as the maritime ports of La Rochelle, Lorient and Brest as well as south of England and Ireland.

